# Prioritization of antimicrobial targets by CRISPR-based oligo recombineering

**DOI:** 10.1101/2021.02.04.429737

**Authors:** HJ. Benns, M. Storch, J. Falco, FR. Fisher, E. Alves, CJ. Wincott, J. Baum, GS. Baldwin, E. Weerapana, EW. Tate, MA. Child

## Abstract

Nucleophilic amino acids are important in covalent drug development yet underutilized as antimicrobial targets. Over recent years, several chemoproteomic technologies have been developed to mine chemically-accessible residues via their intrinsic reactivity toward electrophilic probes. However, these approaches cannot discern which reactive sites contribute to protein function and should therefore be prioritized for drug discovery. To address this, we have developed a CRISPR-based Oligo Recombineering (CORe) platform to systematically prioritize reactive amino acids according to their contribution to protein function. Our approach directly couples protein sequence and function with biological fitness. Here, we profile the reactivity of >1,000 cysteines on ~700 proteins in the eukaryotic pathogen *Toxoplasma gondii* and prioritize functional sites using CORe. We competitively compared the fitness effect of 370 codon switches at 74 cysteines and identify functional sites in a diverse range of proteins. In our proof of concept, CORe performed >800 times faster than a standard genetic workflow. Reactive cysteines decorating the ribosome were found to be critical for parasite growth, with subsequent target-based screening validating the apicomplexan translation machinery as a target for covalent ligand development. CORe is system-agnostic, and supports expedient identification, functional prioritization, and rational targeting of reactive sites in a wide range of organisms and diseases.

## Main text

Electrophilic small molecules that engage protein-encoded amino acid nucleophiles are resurgent in drug discovery as versatile chemical probes and therapeutic agents^1,2^. As a result, considerable efforts are devoted to the development of chemical proteomic technologies termed ‘reactivity-based profiling’ (RBP) for the identification of nucleophilic sites^3^. One such technology, isotopic tandem-orthogonal activity-based protein profiling (isoTOP-ABPP)^4^, has evolved into the standard method for proteome-wide profiling of intrinsic amino acid reactivity. Central to isoTOP-ABPP is the use of quantitative mass spectrometry to measure the extent of protein labelling with a highly-reactive electrophilic probe. Initially applied to rank the reactivity of cysteines in the human proteome using an iodoacetamide probe^4^, isoTOP-ABPP has since been expanded to other amino acid types including lysine^5^, methionine^6^ and tyrosine^7^. Moreover, this method has been successfully adapted for competitive screening of covalent fragments^5,8–10^, enabling identification of sites that can be pursued in fragment-based ligand discovery (FBLD) programs for so-called “inverse drug discovery”^11^.

Despite advances in RBP, the prioritization of reactive or ligandable amino acids as targets following their proteomic identification remains biased; target selection is typically based on the availability of existing functional information or assays for the associated protein class. This inevitably leads to proteins with untapped therapeutic value being overlooked^12^. Following proteomic identification, genetic approaches for functional interrogation of reactive sites are low-throughput and often involve a degree of serendipity. The ability to efficiently interrogate individual reactive amino acids across the proteome at high throughput would expand our understanding of protein sequence-function relationships in complex biological systems, and solve one of the grand challenges of universal inverse drug discovery.

Over recent years, several multiplexed ‘recombineering’ screens (e.g. MAGE, CRMAGE, CREATE) have been developed to simultaneously map the phenotypic effects of thousands of amino acid substitutions across genomes^13–15^. Typically restricted to prokaryotic systems, these platforms monitor the allelic frequency of amino acid mutants in a population over a period of selective pressure or growth, enabling the identification of substitutions that impact cellular fitness. However, these methods indirectly estimate mutant frequency, limiting their ability to probe sequence-function relationships. Other technologies are available that overcome these limitations by direct sequencing of the modified chromosomal loci^16–18^. However, their application has been restricted to single or small panels of targets (e.g. in saturation mutagenesis) and/or a limited range of amino acid substitution types. Therefore, a strategy for direct, quantitative assessment of the contribution of individual amino acids to protein function across a diverse range of genomic loci (such as sites identified by RBP) is needed.

Here, we introduce <u>C</u>RISPR-based <u>O</u>ligo <u>Re</u>combineering (CORe) for proteome-wide assessment of amino acid contribution to protein function in cells. Combined with isoTOP-ABPP, we apply CORe to identify and prioritize reactive cysteines as therapeutic targets of covalent antimicrobials in the eukaryotic pathogen *Toxoplasma gondii.* We reveal the apicomplexan protein translation machinery as an unexpected target for covalent inhibition, and highlight CORe as a general strategy for protein sequence-function studies and the expedient, unbiased prioritization of reactive sites, proteins and biological processes for ligand discovery.

### Cysteine reactivity profiling identifies potential drug targets in *T. gondii*

We sought to establish a platform for the prioritization of druggable sites on protein targets. While the CORe target prioritization platform is conceptually both amino acid - and system-agnostic, for proof-of-concept we focused on electrophile-sensitive cysteines in the apicomplexan parasite *Toxoplasma gondii*. *T. gondii* is an experimentally tractable eukaryotic host-pathogen model^19^ with medical and veterinary importance^20^, and presents resistance to front-line therapeutics^21^, highlighting the need for rapid identification and prioritization of new therapeutic targets.

We began by adapting the ‘Azo’ derivative of the isoTOP-ABPP platform^22^ to *T. gondii*. We profiled the reactivity of protein-associated cysteines in soluble proteome extracts of extracellular *T. gondii* tachyzoites using an iodoacetamide-based probe, IA-alkyne (**Fig. 1a**). After statistical filtering, we identified a total of 1097 cysteines in 691 proteins that were sensitive to IA-alkyne labelling (**Table S1**). Similar to previous studies^4,23,24^, individual cysteines displayed a range of inherent reactivity towards the probe (**Fig. 1b**). Amino acid ‘hyperreactivity’ is an established predictor of functionality in cells^4,5^, and we therefore partitioned cysteines by their respective isotopic ratios into hyper (*R* < 3), medium (*R* = 3-5), and low (*R* > 5) reactivity groups. In total, 130 hyperreactive cysteines were identified in 97 proteins with diverse biological functions (**Fig. S1a**). This includes proteins with known cysteine-based catalytic mechanisms (e.g. thioredoxins), well-characterized parasite proteins for which no functional role has previously been attributed to the identified cysteines (e.g. myosin F), and hypothetical proteins (**Table S2**). Analysis of functional annotations assigned to hyperreactive cysteine-containing genes revealed enrichment of hyperreactive sites enrichment in translation-associated proteins including the ribosome (**Fig. 1c; Table S2, S1a**), which were not correlated with general enrichment of abundant proteins (**Fig. S1b**) and absent from similar datasets obtained from other eukaryotic cell systems^4,23,24^.

**Figure 1.**
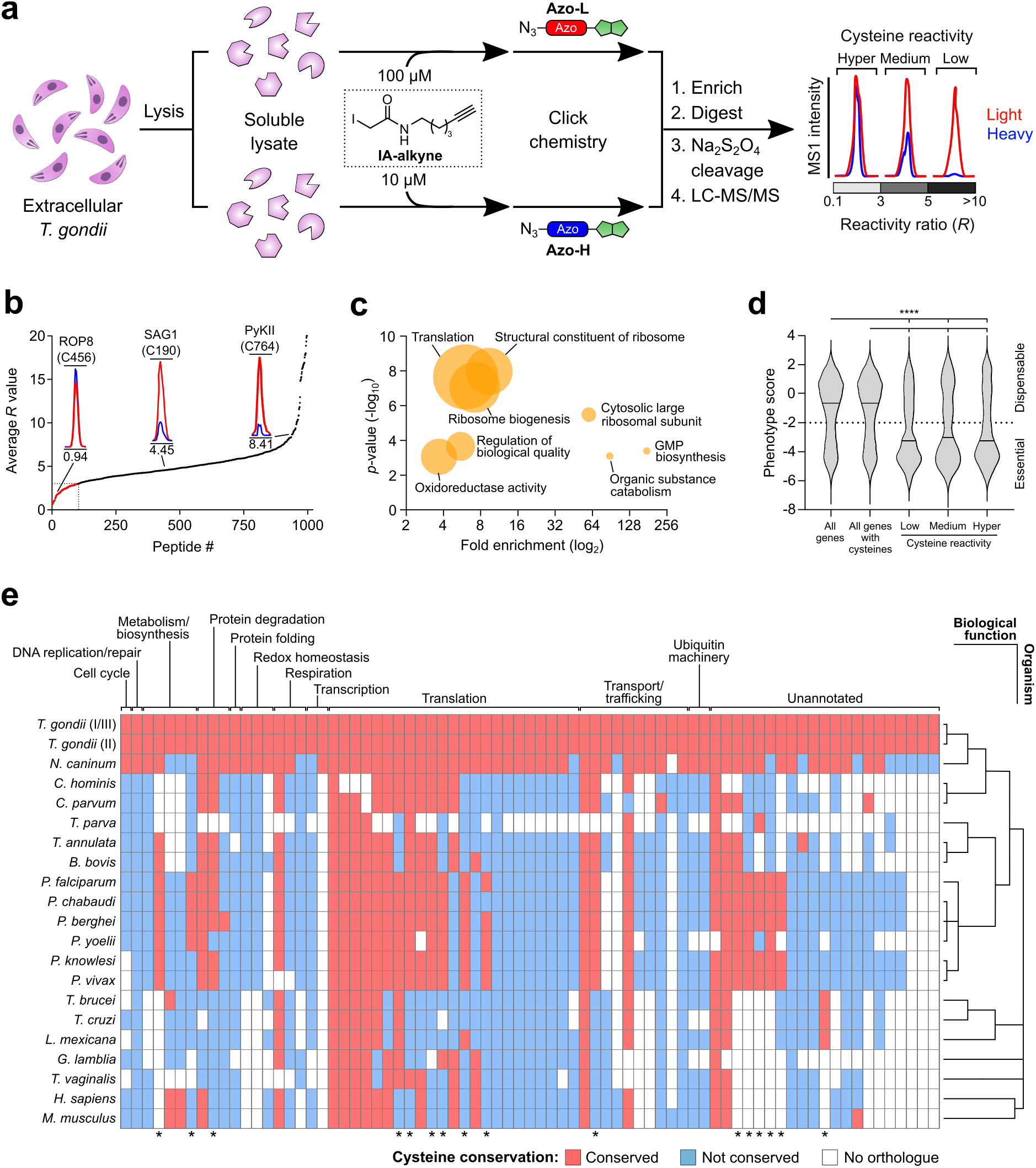
Cysteine reactivity profiling in *T. gondii* reveals enrichment of hyperreactive cysteines in essential and translation-associated proteins. **a.** isoTOP-ABPP workflow for quantifying cysteine reactivity in *T. gondii* parasites. Soluble lysates from extracellular tachyzoites were independently labelled with high (100 μM) and low (10 μM) concentrations of a thiol-reactive IA-alkyne probe. Labelled samples were then click-conjugated to isotopically-differentiated, reductant-cleavable biotin tags (heavy (blue) and light (red) for 10 μM and 100 μM treatment groups, respectively), combined and enriched on streptavidin-immobilized beads. Immobilized proteins were then subject to tandem on-bead trypsin digestion and sodium hydrosulfite treatment, eluting probe-modified peptides for LC-MS/MS analysis. Cysteine reactivity is quantified by *R* values, which represent the differences MS1 peak intensities between the light-and heavy-conjugated proteomes. **b**. Ranked Average *R* values for probe-labelled peptides from two independent experiments (n=2). Representative chromatograms of cysteines within three groups of reactivity (hyper, *R* < 3; medium, *R* = 2-5; low, *R* > 5) are annotated. **c.** Enrichment analysis of functional annotations in hyperreactive cysteine-containing genes relative to the *T. gondii* genome. Fold change is plotted against statistical significance; circle area is proportional to the number of proteins matching with a given term. **d.** Comparative distribution analysis of published phenotype scores^25^ for the *T. gondii* genome with all cysteine- and reactive-containing genes. Essential genes are classified by a score of < −2. **e**. Conservation of hyperreactive cysteines identified in essential *T. gondii* genes across orthologues of eukaryotes. Cysteines are grouped by the predicted function of their associated genes, and organisms by their phylogenetic relationship. Asterisks indicate residues highly conserved in eukaryotic pathogens, but absent in mammalian systems.

We next assessed the association of cysteine reactivity with gene essentiality according to ‘phenotype scores’ from a genome-wide CRISPR knockout screen in *T. gondii* (**Fig. 1d**)^25^. Using a phenotype score threshold of −2 or below as an indicator of gene essentiality, we observed enrichment of indispensable genes in our reactive cysteines dataset relative to all protein-coding genes or protein-coding genes containing at least one cysteine. No difference in the distribution of phenotype scores was observed between the low, medium and hyper reactivity groups. Combined phenotype scoring and bioinformatic analyses identified a focused group of 75 hyperreactive cysteines in 56 essential genes (**Table S2**). Phylogenetic analysis of these targets indicated varying degrees of cysteine conservation across different protein classes and eukaryotes (**Fig. 1e**). Interestingly, several sites appeared to be widely conserved in clinically important pathogens yet absent in the human host, emphasizing the potential for hyperreactive cysteines to be selectively targeted with cysteine-directed drugs. In summary, isoTOP-ABPP captured a unique chemically targetable subset of parasite proteins, highlighting reactivity profiling as a powerful approach to enrich for new potential drug targets in this parasite.

### CORe platform rationale and design

To systematically interrogate cysteines identified by isoTOP-ABPP, we designed a methodology to prioritize individual sites based on demonstrated contribution to protein function in cells. We refer to our approach as <u>C</u>RISPR-based <u>O</u>ligo <u>Re</u>combineering (CORe) (**Fig. 2a)**. The underlying principle of CORe is that for essential genes there is direct relationship between the molecular function of the encoded protein and cellular fitness; mutations that perturb protein function will similarly impact cellular fitness. We hypothesized that by comparing the fitness of wild-type (WT) and cysteine mutants for a specific reactive cysteine-containing gene product, the functional contribution of the target residue can be assessed within the sequence context of the protein. This is achieved through site-specific integration of different mutations using a CRISPR-Cas9-based homology-directed repair (HDR) strategy, and subsequent quantitative comparison of the fitness of the resulting reactive site mutant(s) to WT and knockout (KO) controls.

**Figure 2.**
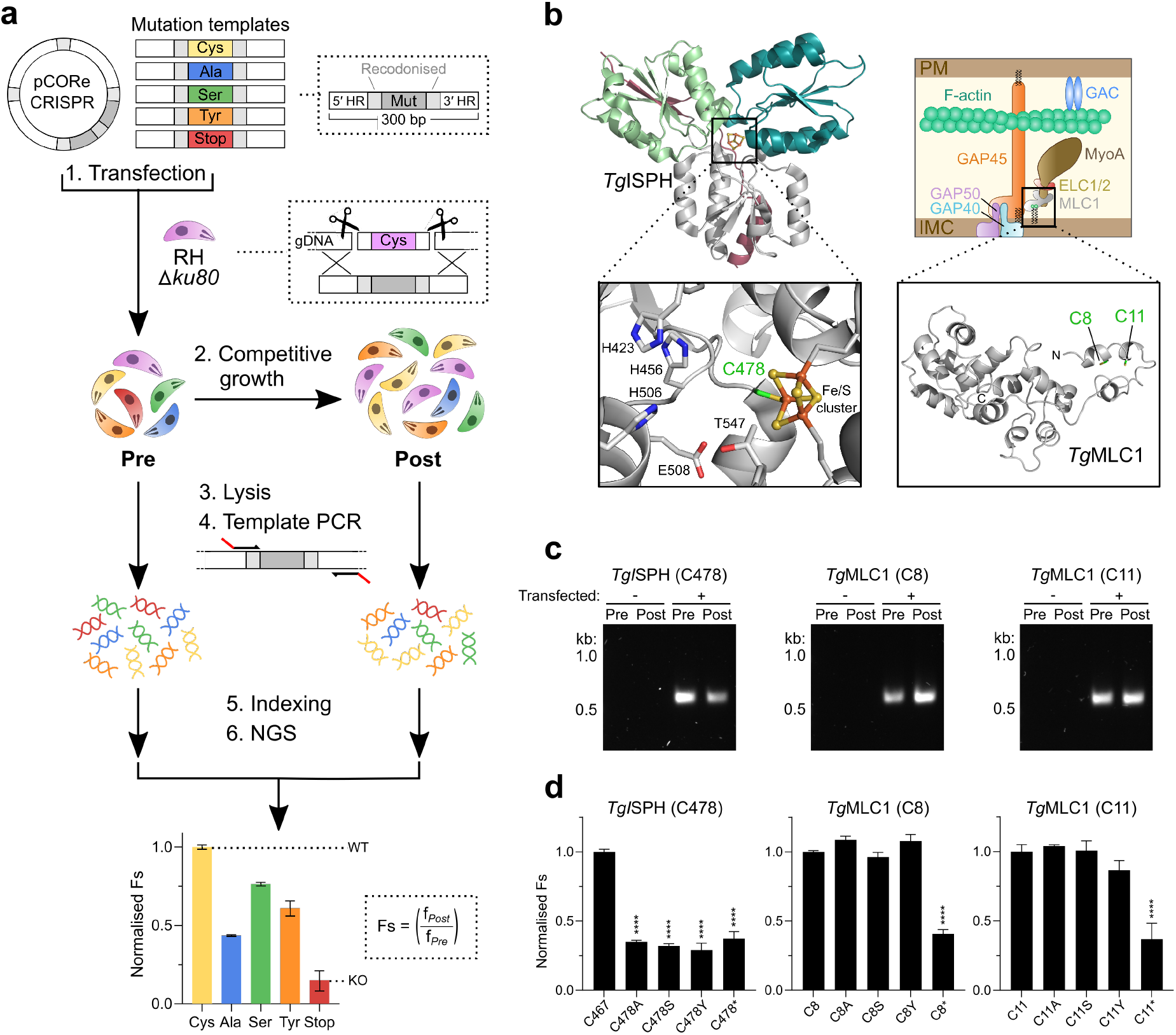
CORe discriminates between essential and non-essential reactive sites. **a.** Workflow of CORe for functional interrogation of hyperreactive cysteines in *T. gondii*. A single pCORe CRISPR plasmid is co-transfected into *T. gondii* parasites with a panel of linear double-stranded donor templates that encode different codon switches (a recodonized cysteine codon, alanine, serine, tyrosine and stop codon). Each plasmid encodes Cas9 nuclease and two gRNA cassettes that direct Cas9 to induce double-stranded breaks (DSBs) at sites 5′ and 3′ of a target cysteine codon. This promotes integration of templates at the excised genomic locus via homology directed repair (HDR), substituting the endogenous cysteine for a given mutation. To increase the efficiency of HDR, a cell line deficient of NHEJ-based DNA repair is used (RHΔ*ku80*)^45^. Genomic DNA from the transfected parasite population is extracted before (‘Pre’) and after (‘Post’) competitive lytic growth. For each time point, specific amplicons are generated by targeting primers to regions of recodonized sequence within the templates. The abundance of each mutation is quantified by next-generation sequencing (NGS). The read frequency of each mutant in ‘Post’ (*f*_Post_) is normalized to ‘Pre’ (*f*_Pre_) to determine fitness scores (Fs) that reflect the viability of parasites following amino acid substitution. Fs values for the amino acid substitutions are compared against the synonymous recodonized cysteine (wildtype) and stop codon (knockout) mutations to identify deleterious mutations (i.e. functional cysteines). **b.** Structural models of CORe targets *Tg*ISPH (left) and *Tg*MLC1 (right). Insets show the positions of their associated target cysteines. **c**. Amplicons generated following mutation of *Tg*ISPH (C478) and *Tg*MLC1 (C8/C11). **d.** Histograms showing Fs values for cysteine mutants of *Tg*ISPH (C478) and *Tg*MLC1 (C8/C11), normalized to the recodonized cysteine control. Data represent mean ±s.d. values for three independent experiment (n=3). Statistical significance was determined by one-way analysis of variance. \*\*\*\**p* < 0.0001.

For functional interrogation of hyperreactive cysteines in *T. gondii*, we selected five mutation types; a recodonized cysteine (synonymous replacement of the target cysteine; WT), a stop codon (for disruption of the target gene; KO^26^), and three distinct amino acid substitutions: alanine, serine or tyrosine. While alanine and serine are commonly used in mutagenesis studies, tyrosine was included to probe sites that could participate in protein-protein interactions (PPIs). Meta-analysis of PPI mutation datasets obtained from cancer studies^27^ revealed that that tyrosine is the most frequent cysteine substitution that causes destabilization at PPI interfaces (**Table S3**). We therefore reasoned that a destabilizing tyrosine mutation may facilitate the identification of cysteine-dependent PPI hotspots, while acknowledging the caveat that a large aromatic substitution may affect protein function via folding defects. Full details on the optimization of CORe are provided in ‘Methods’.

### CORe prioritises cysteine targets according to contribution to protein function in cells

We first trialed CORe against reactive cysteines in two targets: *Tg*ISPH (TGGT1_227420) and *Tg*MLC1 (TGGT1_257680) (**Fig. 2b**). ISPH (also known as ‘LytB’) is an oxidoreductase essential for isoprenoid biosynthesis and an established antimicrobial drug target^28–30^. The reactive cysteine identified in *Tg*ISPH contributes to an iron-sulfur cluster that is required for enzyme catalysis, and therefore any substitutions at this site are expected to be deleterious. *Tg*MLC1 is part of the glideosome complex required for parasite motility and host-cell invasion^31^ and contains two N-terminal reactive cysteines that are known to be *S*-acylated^32,33^ yet are non-essential for *Tg*MLC1 function^34^. We therefore hypothesized that substitutions at this site would not affect parasite fitness. We applied CORe to these targets; integration-specific amplicons were successfully generated (**Fig. 2c**), and NGS analysis confirmed our expectations with high biological reproducibility. Any substitution of the *Tg*ISPH-associated cysteine negatively impacted parasite fitness, with all three mutations being analogous to disruption of the gene following integration of the stop codon (**Fig. 2d**). In agreement with published data, the cysteines on *Tg*MLC1 were permissive to all mutations indicating that these residues (and their post-translational modification) do not contribute to the essential component of this protein’s function (**Fig. 2d**). We next benchmarked a standard genetic analysis workflow against which CORe could be compared. For this purpose, we selected a hyperreactive cysteine associated with a hypothetical protein (TGGT1_258070), generated an inducible KO (iKO) line using the DiCre system (RH *Tg*Hypo^iKO^) (**Fig. S2a**), and confirmed the expected genomic rearrangement by PCR, expression of the protein by Western blot, and localization by immunofluorescence microscopy (**Fig. S2b, d and e**). Treatment of RH *Tg*Hypo^iKO^ parasites with rapamycin resulted in efficient gene excision (**Fig. S2c**), and in agreement with the gene’s phenotype score (−5.24) plaque assay confirmed that knockout parasites were not viable (**Fig. S2f and g**). We then sought to assess the contribution of the reactive cysteine to protein function by genetic complementation in the genetic background of this iKO. Despite repeated attempts we were unable to complement for the loss of this gene, precluding functional interrogation of the associated cysteine. In this instance, the standard approach took ~12 months.

We proceeded to apply CORe to our complete set of essential, hyperreactive cysteine-containing genes. Construction of 59 CRISPR plasmids was accomplished in five days using a linker-based DNA assembly strategy (**Fig. S3a and b**), followed by parasite transfection and competitive lytic growth (eight days), integration-specific amplicon production (achieving 100% coverage for our target cysteines, **Fig. S4**), NGS library construction (seven days) and Illumina NextSeq processing (seven days). The entire CORe workflow took approximately one month to complete for 74 reactive cysteine targets (>800× faster than the standard workflow on a per-gene basis), with the final dataset indicating exceptional reproducibility across independent biological replicates. These data are summarized on **Figures 3a and S5**. For ~90% of the target cysteines (66/74), the integration of the premature stop codon resulted a significant (*p* < 0.05) reduction in parasite fitness (**Fig. 3b**). No deleterious growth phenotype was detected for stop codon mutants in eight targets. This may reflect the proximity of the mutagenized cysteine to the protein C terminus, as these proteins likely retain functional domains (**Fig. S6a**). Interestingly, the relative magnitude of the effect of integrating the stop codon did not correlate with published gene phenotype scores (**Fig. S6b**).

**Figure 3.**
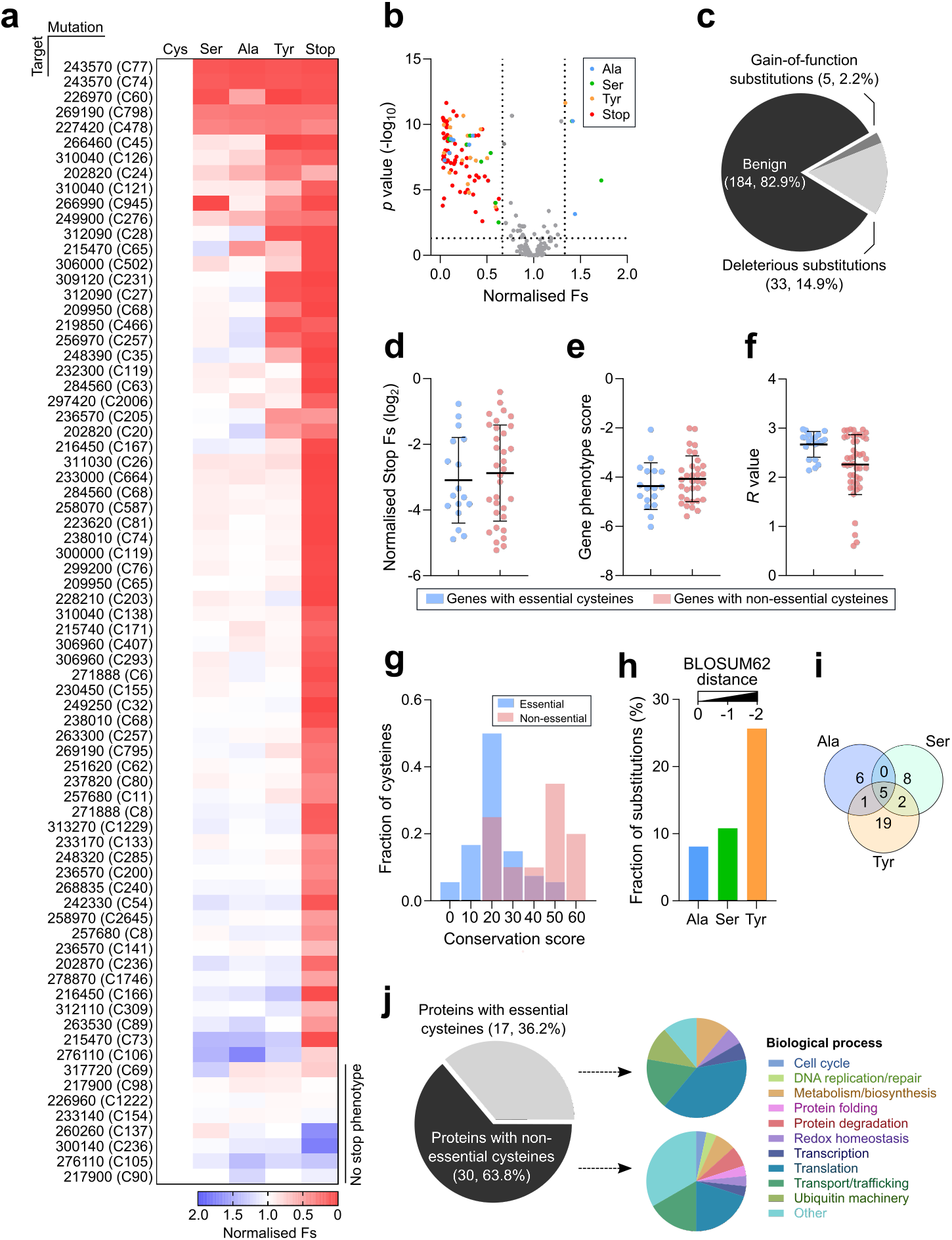
CORe prioritises apicomplexan protein translation as a target for covalent inhibition. **a**. Heatmap showing normalized Fs values for all target cysteines and mutation types ordered by the mutation sensitivity of the cysteines (high to low, top to bottom). **b.** Volcano plot showing the normalized Fs values of each cysteine mutation and significance against the recodonized cysteine control as determined by one-way analysis of variance. Significant mutations (*p* < 0.05) with Fs values < 0.66 and > 1.33 are coloured. **c.** Proportion of amino acid substitutions causing deleterious or gain-of-function phenotypes. **d-f.** Distribution of normalized stop codon Fs values (**d**), phenotype scores (**e**) or isoTOP-ABPP *R* values (**f**) between proteins containing at least one essential or non-essential cysteine. **g.** Frequency distribution of conservation scores assigned to essential and non-essential cysteines across 20 eukaryotic organisms; higher scores indicate wider conservation across the analyzed species. **h.** Fraction of deleterious amino acid substitutions for each mutation type. The BLOSUM62 distance scores for each substitution are annotated and organized by increasing distance from the native cysteine residue (left to right)^35^. **i.** Overlap of cysteines with deleterious alanine, serine and/or tyrosine substitutions. **j**. Proportion and functional annotations of proteins containing essential and non-essential cysteines.

Analysis capturing aspects of both the magnitude and statistical significance of the effect of each individual substitution provided a straightforward route to identify robustly essential cysteines, and prioritize target sites according to their contribution protein function in live cells (**Fig. 3b and Table S4)**. The majority of substitutions were benign (~83%, 184/222), with only a small fraction of reactive cysteines measurably contributing to the function of the protein (~17%, 38/222) (**Fig. 3c**). Unexpectedly, CORe identified gain- as well as loss-of-function mutations. Illustrating the challenge of selecting targets in the absence of an approach such as CORe, there was no association between the essentiality of a reactive cysteine and the effect of stop codon integration (**Fig. 3d**), the phenotype score of the associated gene (**Fig. 3e**), or the reactivity of the cysteine itself (**Fig. 3f**). The challenge of target selection is exemplified by the reactive cysteine originally chosen for validation via the standard genetic workflow (**Fig. S2**). For this hypothetical protein, CORe indicated that the reactive cysteine does not contribute to protein function (**Fig. 3a, S5**). Addressing the relationship between cysteine “essentiality” and function, we compared the extent of conservation for essential and non-essential cysteines according to ‘conservation scores’ (**Fig. 3g and Table S2**). While non-essential cysteines appeared to be normally distributed across the analyzed species, essential cysteines displayed a bimodal distribution with higher scores. This indicated that conservation should not be taken as the sole predictor of function.

Integration of three different amino acid substitutions enabled deeper interrogation of each site, and an increased appreciation of functionally disruptive biochemistry (**Fig. 3h**). In agreement with anticipated evolutionary mutational tolerance, the greater the BLOSUM62 matrix distance^35^ between the individual mutation and cysteine, the more likely the mutation affected the function of the associated protein. This supports a degree of functional buffering or resistance against gradual evolutionary change of protein function as a result of changes in protein sequence, with a range of tolerance observed for each individual cysteine (**Fig. 3i**). Finally, to identify biological processes with potential sensitivity to cysteine-reactive covalent small molecules, we performed an enrichment analysis of functional annotations assigned to proteins containing essential or non-essential cysteines (as defined by CORe) (**Fig. 3j**). The breakdown for the two groups was distinct, with translation annotation enriched in essential cysteine-containing genes. Translation was therefore prioritized for further targeting.

### Protein translation in *Plasmodium falciparum* is sensitive to cysteine-based covalent inhibition

We undertook an in-depth analysis of CORe-prioritized reactive cysteines present on proteins associated with translation. The majority of these sites (9/10) were encoded in proteins decorating the surface of the cytoplasmic 80S ribosome (**Fig 4a and b**). Translation has track record as a therapeutic target, including in the related apicomplexan parasite and etiologic agent of malaria, *Plasmodium falciparum*^36,37^. This parasite remains the cause of significant mortality and morbidity worldwide, and due to existing and emergent drug resistance there is a constant demand for new therapeutic targets and modalities. Our conservation analyses indicated that the majority of essential translation-associated cysteines in *T. gondii* were conserved in *P. falciparum*. Interestingly, not all were conserved in humans, indicating the possibility of parasite specific functions that could be therapeutically targeted (**Fig. 4a and c**). For this subset of parasite-specific cysteines, the tyrosine was the only observed deleterious substitution (**Fig. 4d**), highlighting potential association for one or more of these sites with PPIs. We took advantage of a recently established *in vitro* translation (IVT) assay^38^ to test the sensitivity of both *P. falciparum* and human translation to covalent inhibition with the promiscuous cysteine alkylating molecule, iodoacetamide. Excitingly parasite translation, but not human translation, was uniquely sensitive to inhibition by iodoacetamide (**Fig. 4e**), confirming this as a new potential therapeutic modality for this biological process, and paving the way for future covalent fragment-based ligand discovery.

**Figure 4.**
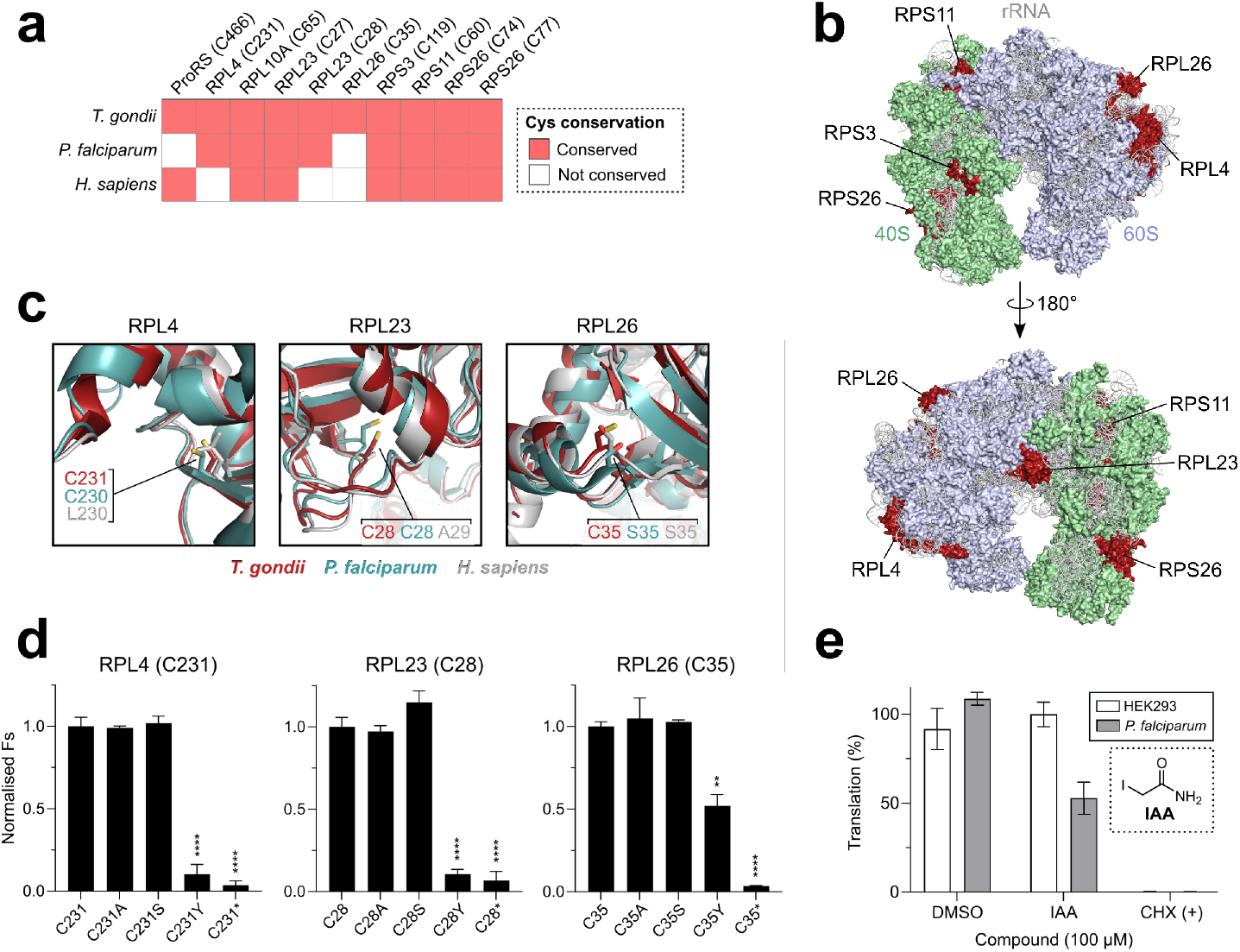
The apicomplexan translation machinery contains unique essential cysteines and is perturbed by covalent modification. **a.** Conservation of essential cysteines identified in translation-associated proteins of *T. gondii* in orthologues of *P. falciparum* and *H. sapiens*. **b.** Front (top) and rear (bottom) views of the cytoplasmic *T. gondii* 80S ribosome (PDB 5XXU/5XXB). Ribosomal subunits, RNA and proteins containing essential cysteines are colored and annotated. **c.** Structural alignment of selected ribosomal proteins (RPL4, RPL23 and RPL26) with orthologues in the *P. falciparum* 80S (PDB 6OKK/3J79) and *H. sapiens* 80S (PDB 4UG0) ribosomes. The essential cysteine residues and their positional equivalents in *P. falciparum* and *H. sapiens* are represented in stick form and annotated. **d.** CORe mutational profiles for RPL4 (C231), RPL23 (C28) and RPL26 (C35). Data shows mean ±s.d. Fs values for each mutation type, normalized to the recodonized cysteine control. **e.** Translational output of *P. falciparum* trophozoite and HEK293 cell lysates following treatment with 100 μM iodoacetamide (IAA). Protein translation was measured using a luciferase-based *in vitro* translation (IVT) assay^38^. Cycloheximide (12 nM) and DMSO treatments were used as positive and negative controls, respectively. Experiments were performed in biological (CORe, n=3) and technical (IVT) triplicate. Statistical significance was determined at n=3 by one-way analysis of variance. \*\*\*\**p* < 0.0001, \*\**p* < 0.01.

## Discussion

In recent years the scope, scale, and speed with which chemically reactive amino acids can be profiled has accelerated dramatically; chemoselective probes are now available for cysteine^4^, serine^39^, lysine^5^, methionine^6^, tyrosine^7^, aspartate/glutamate^9,40^, tryptophan^41^, and histidine^42^. Supporting this, advances in mass-spectrometry have significantly expanded the number and rate at which individual reactive sites can be profiled^43,44^, and subsequently exploited by electrophilic drug hunters. CORe provides a technology bridge between unbiased proteomic profiling of reactive sites and protein sequence-function relationships, and will be a valuable tool for proteome engineers alongside other methods including MAGE, CRMAGE and CREATE. CORe provides a simple strategy to interrogate any individual amino acid, directly assessing its contribution to protein function, which we anticipate will prove as useful as alanine scanning in traditional protein structure-function studies. While *T. gondii* was used for proof-of-concept studies with reactive cysteines, CORe is both system agnostic and amino acid agnostic, with one exciting future application being the systematic profiling of all PTMs of a given class, e.g. sites of phosphorylation or *N*-myristoylation. Many advances in the target identification and validation sphere are currently achieved retrospectively following identification of a suitable ligand. In contrast, global amino acid reactivity profiling combined with CORe supports prospective strategies in target identification and validation campaigns. Our approach enables the critical concept of prioritization to be used to promote protein targets and targetable biological processes into screening platforms where an identified prospectively druggable site is already proven to contribute to protein function in intact cell systems. As such, combined with reactivity profiling, CORe has the potential to focus drug discovery pipelines on functional sites on identified targets, accelerating the discovery of targets and next-generation small molecule therapeutics. The translation of our findings to the related malaria parasite *P. falciparum* provides the first evidence for this potential, with covalent inhibition of apicomplexan parasite translation apparatus being a tantalizing modality for new broad-spectrum antimicrobials.

## Materials and methods

### General

Unless otherwise stated, all reagents were provided by Sigma. All primers/oligonucleotides and synthetic DNA used in this study are listed in Tables S5 and S6, respectively. The IA-alkyne probe, Azo-L and Azo-H tags were synthesized as previously described^1,2^.

### Cell culture and parasite isolation

RH strain *T. gondii* tachyzoites were cultured by serial passage on confluent monolayers of human foreskin fibroblasts (HFF-1 ATCC® SCRC-1041™). HFFs were grown at 37°C and 5% CO_2_ in Dulbecco’s Modified Eagle’s Medium (DMEM) supplemented with 10% (v/v) heat-inactivated foetal bovine serum (FBS), 100 μg/ml penicillin/streptomycin and 2 mM L-glutamine. Unless otherwise stated, parasites were harvested for assays or transfection via mechanical syringe lysis of heavily infected HFFs through a 25-gauge needle.

Highly synchronized 3D7 strain *P. falciparum* asexual parasites were cultured in RPMI-1640 medium supplemented with 0.5% (w/v) AlbuMAX™ II (Life Technologies), 50 μg/ml hypoxanthine, 25 μg/l gentamycin and 0.3 mg/ml L-glutamine. Parasites were routinely cultured at 37°C and 5% CO_2_/3% O_2_ with 2% hematocrit blood (NHS UK Blood Transfusion Service). Media was exchanged daily until the culture reached 10-20% parasitemia with predominantly late trophozoites and early schizonts. Infected red blood cells (RBCs) were isolated by centrifugation (800 × g, 5 min) and lysed in RBC lysis buffer (45 mM HEPES pH 7.45, 100 mM potassium acetate, 1.5 mM magnesium acetate, 2 mM DTT and 0.075% (w/v) saponin) for 10 min at room temperature. The lysed RBCs were then centrifuged (2,800 × g and 4°C, 10 min), and the resulting parasite pellet was suspended in cell lysis buffer (45 mM HEPES pH 7.45, 100 mM potassium acetate, 1.5 mM magnesium acetate, 2 mM DTT). This step was repeated until all RBC debris was removed.

HEK 293F cells were cultured in FreeStyle™ 293 Expression Medium (Life Technologies) at 37°C and 5% CO_2_. Cells were harvested at a density of ~2×10^6^/ml by centrifugation (1000 × g for 10 min at 4°C) and washed once in cell lysis buffer supplemented with 20U of human placental RNase inhibitor and cOmplete™ EDTA-free Protease Inhibitor Cocktail (Roche) prior to processing lysates. All parasite and host cell strains were confirmed negative for the presence of *Mycoplasma* contamination by PCR.

### Plasmid design and construction

To construct pG140::*Tg*Hypo-3×HA, a recodonized *TgHypo* cDNA sequence fused to a C-terminal 3×HA tag was synthesized by GeneArt (Life Technologies). This fragment was cloned into the *Bam*HI and *Hind*III sites of a modified version of the parental plasmid p5RT70loxPKillerRedloxPYFP-HX^3^, in which the *TUB8* promoter had been deleted using the Q5 Site-Directed Mutagenesis Kit (NEB) protocol with primers P1/P2. Next, fragments encompassing the *TgHypo* 5’ or 3’ UTR were PCR amplified from genomic DNA of RHdiCreΔ*ku80*Δ*hxgprt* parasites using primers P3/P4 and P5/P6, respectively. The 5’ UTR fragment was cloned into the *Nar*I site of the intermediate plasmid, followed by the 3’ UTR fragment at the *Sac*I site to generate pG140::*Tg*Hypo-3×HA.

To construct pSAG1::Cas9-U6::sg*Tg*Hypo(×2), Cas9 sgRNA sequences targeting the *TgHypo* 5’ or 3’ UTR were first selected using the Eukaryotic Pathogen gRNA Design Tool (EuPaGDT)^4^. Two single gRNA vectors containing either the 5’ or 3’ UTR-targeting gRNA were then generated using the pSAG1::Cas9-U6::sgUPRT plasmid as a backbone (Addgene #54467)^5^. Here, the parental UPRT-targeting gRNA was replaced with either *Tg*Hypo gRNA using the Q5 Site-Directed Mutagenesis Kit protocol with primers P7/P9 (5’ gRNA) and P8/P9 (3’ gRNA). Next, a fragment encompassing the 5’ gRNA was PCR amplified using primers P10/P11 and Gibson cloned^6^ into the other *Kpn*I and *Xho*I-digested 3’ gRNA plasmid, generating pSAG1::Cas9-U6::sg*Tg*Hypo(×2).

All CORe plasmids were assembled by Biopart Assembly Standard for Indempotent Cloning (BASIC)^7^. To construct the pCORe recipient vector, three DNA parts (a Cas9 nuclease, *hxgprt* selectable marker and an mScarlett counterselection cassette) were generated with flanking BASIC Prefix and Suffix sequences. The Cas9 part was generated via PCR amplification of pCas9/Decoy (Addgene #80324)^8^ using primers P12/P13. The mScarlett part was synthesized by Twist (www.twistbioscience.com). The *hxgprt* part was amplified from pTUB1:YFP-mAID-3HA, DHFR-TS:HXGPRT (Addgene #87259)^9^ using primers P14/P15. Prior to amplification, two internal *Bsa*I sites in the DHFR UTRs of the *hxgprt* cassette were removed using the Q5 Site-Directed Mutagenesis Kit with primers P16/P17 and P18/P19. The resulting DNA parts were cloned into an ampR-p15A backbone in a four-part BASIC reaction, forming pCORe. All BASIC linkers used in the assemblies were synthesized by Biolegio and are listed in Table S6.

### Transfections

All transfections were performed by electroporation using an Amaxa 4D-Nucleofector (Lonza) with program ‘F1-115’. Transfections were carried out using freshly harvested extracellular tachyzoites in P3 buffer (5 mM KCl, 15 mM MgCl_2_, 120 mM Na_2_HPO_4_/NaH_2_PO_4_ pH 7.2, 50 mM D-mannitol).

### Stable parasite line generation

To generate the inducible knockout strain for *Tg*Hypo (here referred to as RH *Tg*Hypo^iKO^), 10 μg of *Sca*I-linearised pG140::*Tg*Hypo-3×HA was co-transfected with 10 μg of pSAG1::Cas9-U6::sg*Tg*Hypo(×2) into 5×10^6^ RHdiCreΔ*ku80*Δ*hxgprt* parasites^10^. Transgenic parasites were selected with 25 μg/μl mycophenolic acid (MPA) and 50 μg/μl xanthine (XAN) 24 hours post-transfection, and individual resistant clones were obtained by limiting dilution. Successful 5’ and 3’ integration of the DNA construct at the endogenous *Tg*Hypo locus was confirmed by PCR using primer P20/P21 and P22/P23, respectively. Disruption of the endogenous *Tg*Hypo locus was confirmed using primers P24/P25. Rapamycin-induced excision of the integrated *Tg*Hypo iKO construct was verified using primers P26/P27.

### Inducible knockout of TgHypo

Confluent HFF monolayers in T25 flasks were infected with ~2-5×10^6^ parasites for 4 hours prior to treatment with 50 nM rapamycin or an equivalent volume of vehicle (DMSO) for 4 hours. After washout, parasites were grown for at least 24 hours prior to PCR or western blot analysis.

### SDS-PAGE and western blot analysis

Extracellular parasites were lysed RIPA buffer (150 mM NaCl, 50 mM Tris-HCl (pH 8.0), 1% Triton X-100, 0.5% sodium deoxycholate, 0.1% SDS, 1 mM EDTA) supplemented with cOmplete™ Protease Inhibitor Cocktail (Roche) for 1 hour on ice. Lysates were then centrifuged (21,000 × g, 30 min at 4°C), and protein concentration in the supernatant was quantified using the Pierce™ BCA Protein Assay Kit (Thermo Scientific). Laemmli buffer was added to the lysate to 1× concentration (2% SDS, 10% glycerol, 5% 2-mercaptoethanol, 0.002% bromophenol blue and 125 mM Tris HCl, pH 6.8) and boiled (95°C, 5 min) before separation by SDS-PAGE on 12% polyacrylamide gels. Thirty micrograms of protein were typically loaded per lane. Proteins were transferred (20 V, 1 min; 23 V, 4 min; 25V; 2 min) to nitrocellulose membranes using an iBlot 2 Dry Blotting System (Invitrogen). Membranes were briefly washed in PBS-T (0.1% Tween-20/PBS), blocked (5% skimmed milk/PBS-T, 1 hour) and incubated with primary antibodies (1% BSA/PBS-T, overnight at 4°C) at the following dilutions: mouse anti-SAG1 (1:1000, Thermo Scientific) and rat anti-HA (1:1000, company, Roche). Following washing (PBS-T, 3×), membranes were incubated with HRP-conjugated secondary antibodies (1:5000, Thermo Scientific) in 1% BSA/PBS-T for 1 hour at room temperature. Protein bands were developed using the ECL™ Western Blotting Detection Reagent (GE Healthcare) and chemiluminescence was visualized using a ChemiDoc MP Imaging System (Bio-Rad).

### Immunofluorescence microscopy

Confluent HFF monolayers grown on glass coverslips were seeded with ~100,000 parasites. Approximately 24 hours post-infection, cells were fixed (4% paraformaldehyde for 15 min at room temperature) permeabilized (0.1% Triton X-100/PBS for 5-10 min) and blocked (3% BSA/PBS for 1 hour at room temperature). Staining was performed for 1 hour with primary antibodies at the following dilutions: rat mouse anti-SAG1 (1:1000, Thermo Scientific), rabbit anti-HA (1:1000, company – check with Fabio) and X anti-Ty1 (1:1000, Baum Lab). Labelled proteins were stained for 1 hour at room temperature using Alexa Fluor 488/594-conjugated goat antibodies (1:2000, Life Technologies). Nuclei were stained using the intercalating DNA dye DAPI at 5 μg/ml. Stained coverslips were mounted onto glass slides using VECTASHIELD® Antifade Mounting Media (Vector Labs) and imaged on a Nikon Ti-E inverted microscope. Images were acquired using an ORCA-Flash 4.0 camera and processed using ImageJ software.

### Plaque formation

Confluent HFF monolayers grown in 6-well plates were seeded with 200-400 parasites. Parasites were allowed to invade overnight prior to treatment with 50 nM rapamycin or DMSO for 4 hours. Following replacement to standard culture medium, plaques were left to form undisturbed for 6-7 days. Monolayers were then fixed with ice-cold methanol for 10 min and stained with crystal violet stain (2.3% crystal violet, 0.1% ammonium oxalate, 20% ethanol) for 2 hours. Plaques were enumerated manually, and statistical significance in plaque counts between rapamycin and DMSO-treated samples were tested using two-tailed unpaired Student’s *t*-tests with unequal variance. The data are presented as mean (±SD) counts.

### Design and optimisation of the CORe platform

The design of the CORe workflow begins with the identification and selection of paired CRISPR guide RNA (gRNA) sequences that target the Cas9 nuclease to sites 5’ and 3’ of a target cysteine codon. As demonstrated in *Caenorhabditis elegans*^11^, we reasoned that a dual gRNA strategy would provide positive selection towards HDR-mediated integration of mutational templates for our essential gene subset, as the lack of repair of two double-strand breaks (DSBs) in an essential gene should be refractory to growth. To test this hypothesis, the frequency of mutants following mutagenesis of an N-terminal proline codon in surface antigen gene1 (*SAG1*) was compared using single or dual gRNAs in combination with single- or double-stranded strand donor repair templates (**Fig. S7a**). These experiments revealed that dual gRNAs in combination with double-stranded templates provided the highest integration efficiency in the absence of any selectable marker. As anticipated in the absence of drug selection, the frequency of mutants was low (**Fig S7b**). The potential negative impact of this upon quantitation of integration events was circumvented through the inclusion of recodonized sequence within the donor template. This allowed for integration-selective priming and therefore generation of PCR amplicons of modified genomic loci for downstream NGS analyses (**Fig. 2a, S7c**). The protein-centric CRISPR guide design tool, CRISPR-TAPE^12^, was used to simplify and accelerate the gRNA identification and selection process for target cysteines. Accommodating the need for high-throughput multiplexed vector construction, BASIC^7^ was adapted to our sequences and used for facile, modular and scalable production of all transfection vectors, with dual gRNA cassettes and Cas9 encoded on the same vector as previously reported (**Fig. S3a**)^8,13^. The RHΔ*ku80* NHEJ-deficient parasite strain was used to further promote HDR^14^.

Donor repair templates were designed to 1) destroy the protospacer adjacent motif (PAM) and/or gRNA seed sequence required for Cas9 targeting and so prevent further modification of the site following integration; 2) provide a recodonized stretch of sequence proximal to the target cysteine for the generation of integration-specific amplicons at mutated sites. Transfection with the dual gRNA vector introduces DSBs 5’ and 3’ of the target cysteine. The excised locus is subsequently repaired using one of the donor templates, producing a mixed mutant pool, which is sampled shortly after transfection for subsequent genomic DNA extraction (‘Pre’ sample) (**Fig. 2a**). For each reactive cysteine candidate, *T. gondii* tachyzoites are co-transfected with a single cysteine-targeting dual gRNA plasmid and all five donor templates for HDR (**Fig. 2a**). The repair templates encoded for either a WT synonymous replacement of the target cysteine, a stop codon, or one of the three amino acid substitution options.

Following transfection, the mixed population of mutants grow competitively, and are sampled for genomic extraction (‘Post’ sample) (**Fig. 2a**). Where the DSB is repaired using the synonymous WT template, parasites are expected to grow normally. In instances where the stop codon template is integrated, the gene coding sequence (CDS) is disrupted, with parasite growth anticipated to be attenuated equivalent to a knockout^15^. After quantitative deep sequencing of integration-specific amplicons encompassing a target cysteine, the frequency of reads for a given mutant in the Post sample (*f*_Post_) is normalized to Pre (*f*_Pre_) to derive fitness scores (Fs) that reflect the viability of parasites during competitive lytic growth. The Fs’ for the amino acid mutants are benchmarked against the synonymous WT and stop codon mutants. This provides a quantitative assessment of the contribution of an individual cysteine to protein function in live cells, using mutant cell fitness as a measurable phenotype and NGS reads as the readout. Multiplexing of CRISPR vector construction with BASIC, 96-well plate-based transfections, and automated an NGS sample preparation workflow enables hundreds of targets to be functionally interrogated in parallel.

### CORe plasmid and template library design and construction

Guide RNAs were searched against the *T. gondii* GT1 genome (release 46; www.toxodb.org) using the ‘position-specific’ function of CRISPR-TAPE (version 1.0.0)^12^. Briefly, gRNAs binding in near proximity of a target cysteine codon were identified by applying a search distance threshold of ±200 nt. For each codon, two gRNAs binding at sites 5’ and 3’ of the residue were then selected. Selection criteria was based on the number of potential off-target sequences, %GC content and the ability to introduce synonymous PAM or guide blocking mutations at the target genomic sequence. gRNAs were synthesized by Twist as a fragment containing a U6 promoter and flanking BASIC Prefix and Suffix sequences, and independently cloned into *Bsa*I sites of a kan^R^-pMB1 storage plasmid, pTwist Kan (High Copy). For each target cysteine, the corresponding 5’ and/or 3’-binding gRNA fragment were subcloned into pCORe in a three-part BASIC reaction, replacing the mScarlett counterselection cassette and generating the pCORe-CRISPR plasmid. The sequences of all gRNA fragments are listed in Table S6.

Donor templates for mutation of target cysteines were synthesized as 300 bp double-stranded fragments by Twist. For the *SAG1* experiments, 70 bp single-stranded oligonucleotides (P28-P27) were used and hybridized to generate double-stranded templates. For each cysteine codon, five templates were designed to incorporate single unique mutations; a recodonized cysteine codon, alanine, serine, tyrosine or a stop codon. Mutation sites were flanked by regions of synonymous recodonized sequence to (1) enable specific detection of cysteine mutants by PCR, and (2) introduce blocking mutations at the PAM and/or gRNA seed sequence to prevent re-excision of modified genomic loci. Recodonisation was avoided or minimised at intron-exon junctions to avoid interference with mRNA splicing. Homology regions were incorporated on either end of templates to promote genomic integration of mutational templates by HDR. The sequences of all mutational templates are listed in Table S6.

### CORe mutagenesis screens

Transfections were carried out in 16-well Nucleocuvette™ strips using the Amaxa 4D-Nucleofector X-Unit (Lonza). For the optimized CORe screen, 7 μg of pCORe-CRISPR and 0.2 μg of each of the five corresponding mutational templates (equivalent to a ~1:5 plasmid-to-template molar ratio) were co-transfected into 1×10^6^ RHΔ*ku80*Δ*hxgprt* parasites^14^. For the *SAG1* experiments, 6 μg of pCORe-CRISPR and 2 μg of a single template were transfected (~1:100 plasmid-to-template molar ratio). Transfected parasites were expanded in HFF monolayers grown in 24 well plates and allowed to egress naturally three days after infection. Approximately 2×10^6^ of the egressed parasites were used to infect confluent HFF monolayers in 6 well plates, and the remaining parasites (~2×10^6^) were pelleted and frozen for genomic DNA extraction as the initial ‘Pre’ mutant population. Parasites were allowed to egress naturally five days after infection and similarly harvested as the ‘Post’ mutant population. Parasite genomic DNA from frozen cell pellets was extracted using the DNeasy Blood & Tissue Kit (Qiagen) for downstream NGS library preparation.

### Illumina library preparation, sequencing and data analysis

Genomic DNA libraries were prepared similarly to the 16S Metagenomic Sequencing Library Preparation guide (Illumina). Briefly, for each target cysteine, a ~600-800 bp fragment targeting the modified genomic locus was PCR amplified from parasite DNA. For the *SAG1* experiments, the amplicons were designed to encompass the template integration site of both modified and unmodified loci. All primers were designed to include overhanging Illumina adapter sequences and are listed in Table S5 (P32-P181). The resulting amplicon was purified using AMPure XP magnetic beads (Beckman Coulter). Dual indices and sequencing adapters were then ligated to the purified products using the Nextera XT Index Kit (Illumina). Indexed amplicons were then purified using AMPure XP beads, and quantified using the Qubit™ dsDNA HS/BR Assay Kits (Invitrogen), or the QuantiFluor ONE dsDNA System (Promega). Indexed amplicons were pooled at equimolar concentration, and the size and purity of the resulting library was assessed on a TapeStation 2200 with the D1000 ScreenTape System (Agilent). The transfer of reagents used for the purification and indexing of amplicons was performed using acoustic liquid handling (Echo 525, Labcyte). Pooled libraries were sequenced using an Illumina NextSeq 500 75PE Mid Output run with a PhiX spike-in of 10%. Following acquisition, sequencing data were demultiplexed using CASAVA 2.17 and analyzed using the Galaxy web server (www.usegalaxy.org). For each uniquely indexed sample, the sequences were concatenated and separated by each template variant to determine the read counts of the different mutation types. The change in frequency of each mutant variant was calculated by normalizing the percent proportion of reads in the Post population sample to the Pre. The differences in normalized read frequency of the nonsynonymous mutations were statistically tested against the recodonized cysteine mutation by one-way analysis of variance (ANOVA).

### Cysteine labelling and click chemistry

Cell pellets of *T. gondii* RHΔ*ku80*Δ*hxgprt* parasites were lysed by sonication in PBS (pH 7.4) and soluble fractions separated by centrifugation at 3,500 × g for 5 min. Protein concentrations were determined using the DC Protein Assay Kit (Bio-Rad) and a SpectraMax M2e Microplate Reader (Molecular Devices). Proteome samples diluted to 2 mg/ml were treated with 10 or 100 μM IA-alkyne (from 1 mM and 10 mM stocks in DMSO, respectively) and incubated for 1 hour at room temperature with rotation. The labelled proteins were then subject to click chemistry by addition of 100 μM Azo-L or Azo-H, 1 mM TCEP, 100 μM TBTA, and 1 mM CuSO_4_ (final concentrations). Click reactions were incubated for 1 hour at room temperature with shaking. The Azo-L/H-labelled protein samples were then precipitated by adding trichloroacetic acid (TCA) to 10% (v/v) concentration. After overnight storage at −80°C, precipitated proteins were pelleted by centrifugation (15,000 rpm, 10 min), washed 3× with chilled MeOH and resolubilized in 1.2% SDS in PBS by gentle sonication and heating (80°C, 10 min).

### Enrichment and on-bead digestion

Labelled proteome samples were diluted to 0.2% SDS with PBS. The resulting samples were then added to 100 μl of Pierce™ Streptavidin beaded agarose resin (Thermo Scientific) and incubated overnight at 4°C followed by a further 2 hours at room temperature. Protein-bound beads were washed with 1× 0.2% SDS in PBS, 3× PBS and 3× H_2_O before resuspending in 6 M urea in PBS +10 mM DTT and incubating at 65°C for 15 min. Reduced samples were then alkylated by adding iodoacetamide to a final concentration of 20 mM and incubating for 30 min at 37°C with rotation. Samples were diluted 3-fold with PBS and centrifuged (1400 × g, 2 min) to pellet the beads. The beads were resuspended in a mixture of 200 μl of 2 M urea in PBS, 1 mM CaCl_2_ and 2 μg trypsin and incubated overnight at 37°C. The beads were separated from the digest by centrifugation and washed 3× with PBS and 3× H_2_O. Azo-labelled peptides were then cleaved by adding 50 mM sodium hydrosulfite (Na_2_S_2_O_4_) and rotating at room temperature for 1 hour. Eluted peptides were then collected from the supernatant, and Na_2_S_2_O_4_ cleavage was repeated twice more to fractionate the sample. Between each cleavage, the beads were washed with 2× H_2_O and combined with the previous elution. Formic acid was added to the sample to 20% (v/v) concentration before storing at −20°C until mass spectrometry analysis.

### LC/LC-MS/MS analysis, peptide identification and quantification

LC-MS/MS analysis was performed on an LTQ-Orbitrap Discovery mass spectrometer (Thermo Scientific) coupled to an Agilent 1200 Series HPLC. Azo digests were pressure loaded onto 250 μm fused silica desalting columns packed with 4 cm Aqua C18 reverse phase resin (Phenomenex). Peptides were then eluted onto a biphasic column consisting of 100 μm fused silica packed with 10 cm C18 and 4 cm PartiSphere SCX resin (Whatman) following a five-step multidimensional LC/LC-MS/MS protocol (MudPIT)^1^. Each step used a salt push (0%, 50%, 80%, 100%, 100%) followed by an elution gradient of 5-100% Buffer B in Buffer A (Buffer A: 95% H_2_O, 5% MeCN, 0.1% formic acid; Buffer B: 20% H_2_O, 80% MeCN, 0.1% formic acid) at a flow rate of 250 nl/min. Eluted peptides were injected into the mass spectrometer by electrospray ionization (spray voltage set at 2.75 kV). For every MS1 survey scan (400-1800 m/z), 8 data-dependent scans were run for the n^th^ most intense ions with dynamic exclusion enabled.

The generated tandem MS data were searched using the SEQUEST algorithm^16^ against the *T. gondii* database (GT1 proteome), *Toxo*DB (http://toxodb.org/). A static modification of +57.02146 on cysteine was specified to account for alkylation with iodoacetamide. Variable modifications of +456.2849 and +462.2987 were further assigned on cysteine to account for the probe modification with the isotopically light (Azo-L) and heavy (Azo-H) variant of the IA-alkyne-Azo adduct, respectively. Output files from SEQUEST were filtered using DTASelect 2.0. Quantification of isotopic light:heavy ratios was performed using the CIMAGE quantification package as previously described^17^. Overlapping tryptic peptides containing the same labelled cysteine (but different charge states or tryptic termini) were grouped and the median reported as the final light:heavy ratio (*R*). *R* values were averaged across biological replicates and peptides with relative standard deviations of the ≥ 50% *R* value were removed.

### Bioinformatics analysis of reactive cysteine dataset

Functional annotation of reactive cysteine proteins was carried out using BLASTP, Gene Ontology (GO) and InterPro searches within Blast2GO 5 PRO software^18^. Consensus protein sequences were BLASTP searched against the non-redundant (nr) NCBI protein database using an *E*-value cut-off of 10^−6^. GO terms (molecular function, biological process and subcellular localization) were then mapped from the top 20 hits and merged with annotations derived from the InterPro database (www.ebi.ac.uk/interpro). Assignments were further optimized using Annex augmentation^19^.Enrichment of annotations was assessed using a Fisher’s exact test against the *T. gondii* proteome (strain GT1; UniProt Taxonomy ID 507601) at < 0.05 FDR.

For conservation analyses of reactive cysteines, orthologues of the associated protein were identified from orthologue groups classified on OrthoMC^20^. Conservation of a given residue was assessed following BLASTP alignment of the orthologous protein sequence against the *T. gondii* template sequence. Scores were assigned to each alignment based on the presence or absence of a matched cysteine; a score of 3 was assigned to conserved cysteines, 1 for no conservation, and 0 if no protein was identified in the orthologue group for a given species. Conservation scores were determined for each cysteine by summing of the scores across the analyzed species.

### In vitro translation (IVT) assay

Pellets of *P. falciparum* 3D7 or HEK 293F cells were suspended in 1× pellet volume of lysis buffer supplemented with 20U of human placental RNase inhibitor and cOmplete™ EDTA-free Protease Inhibitor Cocktail (Roche). Resuspended parasites were then transferred to a prechilled nitrogen cavitation chamber (Parr Instrument Company) and incubated on ice at 1500 PSI for 60 min. Following release from the chamber, the crude lysate was clarified by differential centrifugation (15 min at 10,000 × g and 4°C, followed by 15 min at 30,000 × g and 4°C). Protein concentration was determined using a NanoDrop (Thermo Scientific) at 280 nm and adjusted to 12 mg/ml prior to storage at −80°C. Prior to performing *in vitro* translation assays, low-bind 384-well plates (Corning) were printed (D300e Digital Dispenser, Tecan) with compounds dissolved in DMSO to be assayed at 0.5% of the total assay volume. Five microlitres of *P. falciparum* clarified lysate was then added to each well, followed by 4.5 μl L-amino acids (each at 200μM in 45 mM HEPES pH 7.45, 100 mM potassium acetate, 1.5 mM magnesium acetate, 2 mM DTT, 20 U human placental RNase inhibitor, 15 μM leupeptin, 1.5 mM ATP, 0.15 mM GTP, 40 U/ml creatine phosphokinase and 4 mM creatine phosphate (Thermo Scientific), 2% (w/w) PEG3000, 1 mM spermidine and 0.5 mM folinic acid) and 0.45 μl of purified red click-beetle luciferase (CBG99) mRNA (1 μg/μl). CBG99 mRNA was transcribed from expression plasmids pH-CBG99-H (for use in *P. falciparum* assays) or pT7CFECBG99 (HEK 293F assays) as previously described^21^. Prepared plates were incubated at 32°C for 1 hour 40 min before adding 10 μl of 45 mM HEPES pH 7.45, 1 mM magnesium chloride, 1 mM ATP, 5 mM DTT, 1% (v/v) Triton-X, 10 mg/ml BSA, 1× Reaction Enhancer (Thermo Scientific), 1 mg/ml D-luciferin (Thermo Scientific) and 0.5 mM cycloheximide. Luminescence was measured across each well using a Tecan M200 Infinite Pro microplate reader heated to 37°C.

### Protein structures and homology modelling

Solved protein structures were downloaded from the RCSB PDB (www.rcsb.org). Homology models were predicted from primary protein sequences using the Phyre^2^ web portal^22^; only models constructed with 100% confidence and ≥ 40% sequence identity across ≥ 70% of the sequence were used. Structural images were generated using PyMOL software (version 2.1.1.; Schrödinger LLC).

### Statistical analysis

Statistical tests were performed using GraphPad Prism 8.0 as described in the individual experimental sections above. *P*-value significance thresholds were set at: **** = *p* < 0.0001, *** = *p* < 0.001, ** = *p* < 0.01 and * = *p* < 0.05. All significant results are annotated with a line and asterisk(s) in the graphs.

### General software

Schematics were created using Adobe Illustrator (version 22.1) and Inkscape (version 0.92.3). Chemical structures were drawn in ChemDraw Professional (version 18.0). PyMOL (version 2.1.1) was used to generate 3D protein structures.

## Acknowledgements

This work was supported by grants BB/M011178/1 from the BBSRC (to HJB, EWT, and MAC) and 202553/Z/16/Z from the Wellcome Trust & Royal Society (to MAC). We would like to acknowledge and thank Ivan Andrew and Laurence Game at the UKRI London Institute of Medical Sciences Genomics Laboratory for providing resources and support that contributed to the research results reported within this paper. We would also like to thank Prof. Matthew Bogyo and Dr. Moritz Treeck for early guidance and support for this work.

## Author contributions

Investigation: HJB, MS, JF, FF, EA and CJW, MAC. Formal analysis: HJB, FF, EW, MAC. Visualization: HJB and MAC. Conceptualization: MAC. Writing—original draft: HJB, EWT and MAC. Writing—review and editing: HJB, MS, FF, EA, CJW, JB, GB, EW, EWT and MAC. Supervision: JB, GB, EW, EWT and MAC. Funding acquisition: JB, GB, EW, EWT and MAC.

## Competing interest declaration

The authors declare no conflict or competing interests.

**Additional information (containing supplementary information line (if any) and corresponding author line),**

## Extended data

**Table S1. Raw isoTOP-ABPP MS data and statistical filtering**

**Table S2: Hyperreactive cysteines and bioinformatics analyses**

**Table S3.** Tyrosine is the most frequent cysteine mutation that causes destabilisation of protein-protein interactions (PPIs) in cancer-associated genes. Shown is the frequency of different amino acid mutations that cause destabilisation of PPIs, where cysteine is endogenous residue. Tyrosine was found to account for the highest proportion (highlighted red). Data obtained from Engin et al.^1^

| Mutation | Count | Frequency (%) |
| --- | --- | --- |
| F | 4 | 14.3 |
| G | 5 | 17.9 |
| R | 5 | 17.9 |
| S | 2 | 7.1 |
| W | 3 | 10.7 |
| Y | 9 | 32.1 |
| **Total** | **28** | **-** |

**Table S4. CORe dataset and essential cysteines**

**Table S5. Primers used in the study**

**Table S6. Synthetic DNA used in the study**

---

**Figure S1.**
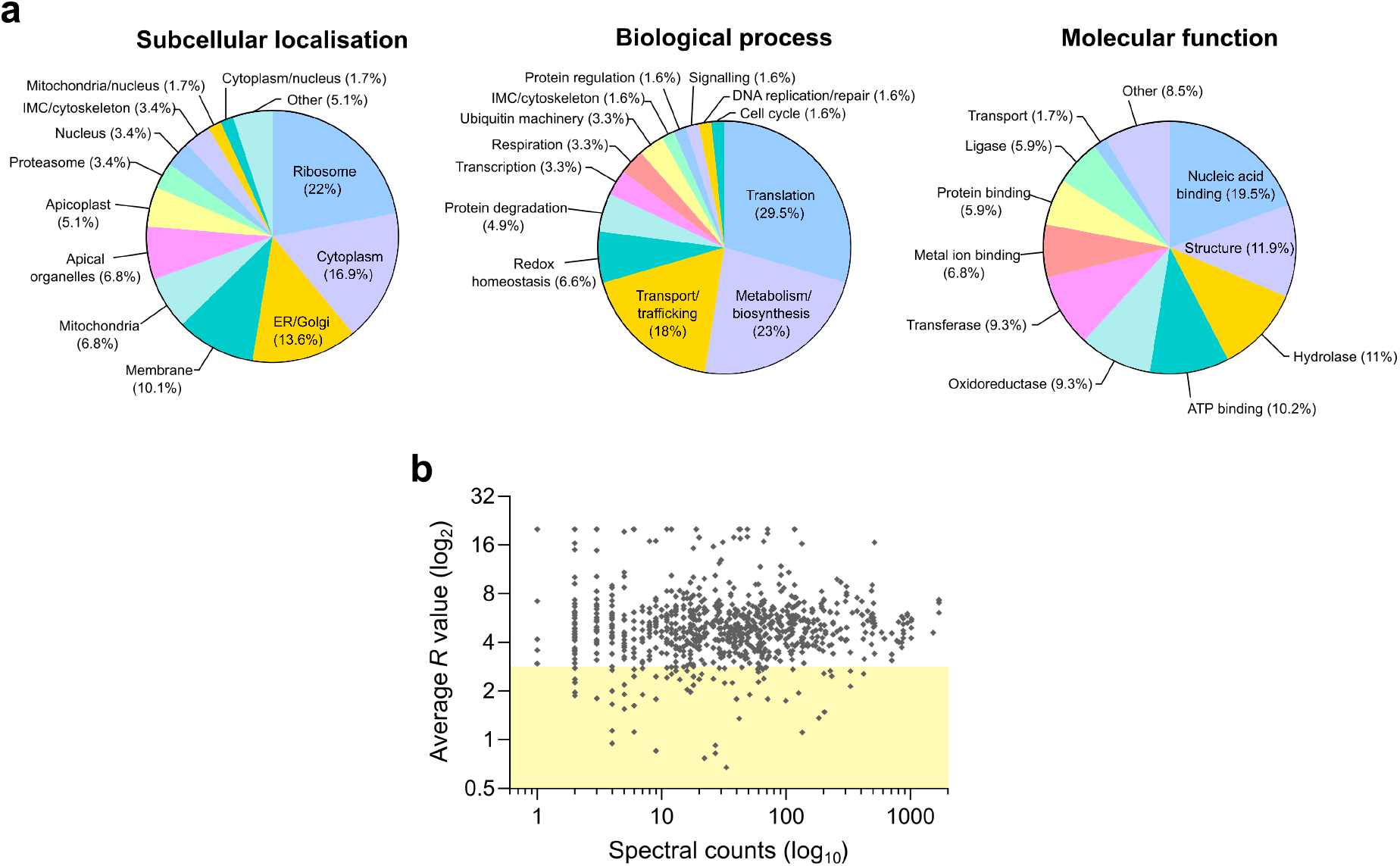
Enrichment hyperreactive cysteines in ribosomal proteins is independent of protein abundance. **a.** Proportions of hyperreactive cysteine-containing genes with functional annotations for three gene ontology categories; subcellular localization, biological process and molecular function. Pie charts depict overrepresentation of naturally abundant proteins, such as ribosome components. **b**. Linear regression analysis of isoTOP-ABPP *R* values against total spectral counts of the associated proteins (a semi-quantiative measure of protein abundance). Spectral counts were obtained from a published proteomic dataset for extracellular *T. gondii* parasites^2^. Note that proteins with low *R* values (< 3, highlighted) span a broad range of spectral counts.

**Figure S2.**
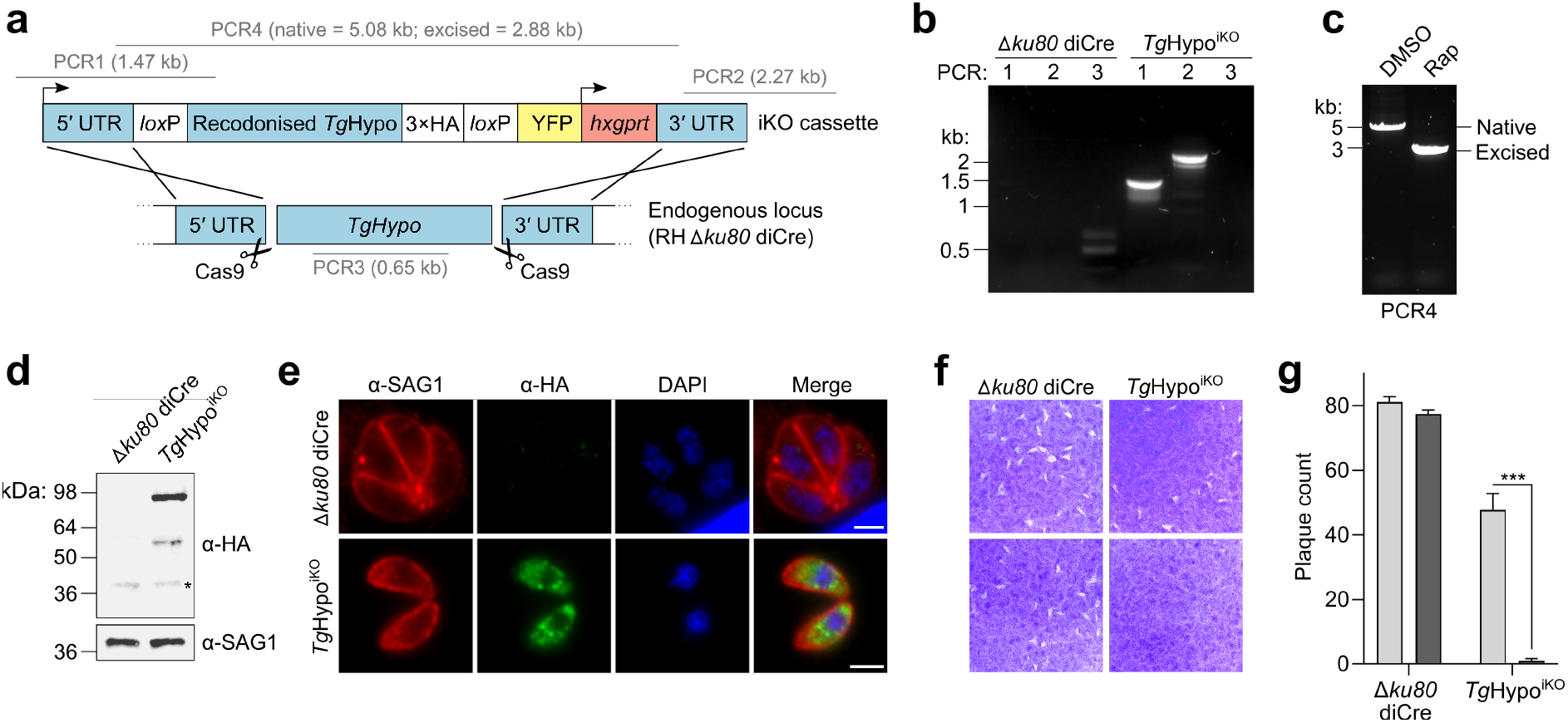
*Tg*Hypo is indispensable for *T. gondii in vitro*. **. a**. Schematic of the CRISPR-based HDR strategy used for generating a *Tg*Hypo inducible knockout line (TgHypo^iKO^) using the diCre system. The predicted sizes of the PCR amplicons used for validating genomic integration and excision of loxP-flanked gene constructs are annotated. **b.** PCR products confirming correct integration of the floxed *Tg*Hypo construct at the 5’ and 3’ UTRs, and loss of the wildtype *Tg*Hypo at its endogenous locus. **c.** Analytical PCR showing complete excision of the floxed *Tg*Hypo-3×HA construct in *Tg*Hypo^iKO^ parasites. **d.** Western blot showing expression of the 3×HA-tagged *Tg*Hypo construct in *Tg*Hypo^iKO^ parasites using an α-HA antibody; equal protein loading was verified using an α-SAG1 antibody. **e.** Immunofluorescence micrographs of *Tg*Hypo^iKO^ parasites following staining with α-HA antibodies, showing correct cytosolic localization of the *Tg*Hypo-3×HA construct. SAG1 and DAPI were used as parasite surface and nuclear markers, respectively. Scale bar = 3 μm. **f.** Representative images of plaques formed on HFF monolayers by the indicated strains in the presence of rapamycin or DMSO. **g.** Plaque counts for each strain determined from (**f**) showing loss of plaquing capacity in *Tg*Hypo^iKO^ parasites upon rapamycin treatment. Data represents three biological replicates (n=3). Statistical significance was determined by one-way analysis of variance. \*\*\**p* < 0.001.

**Figure S3.**
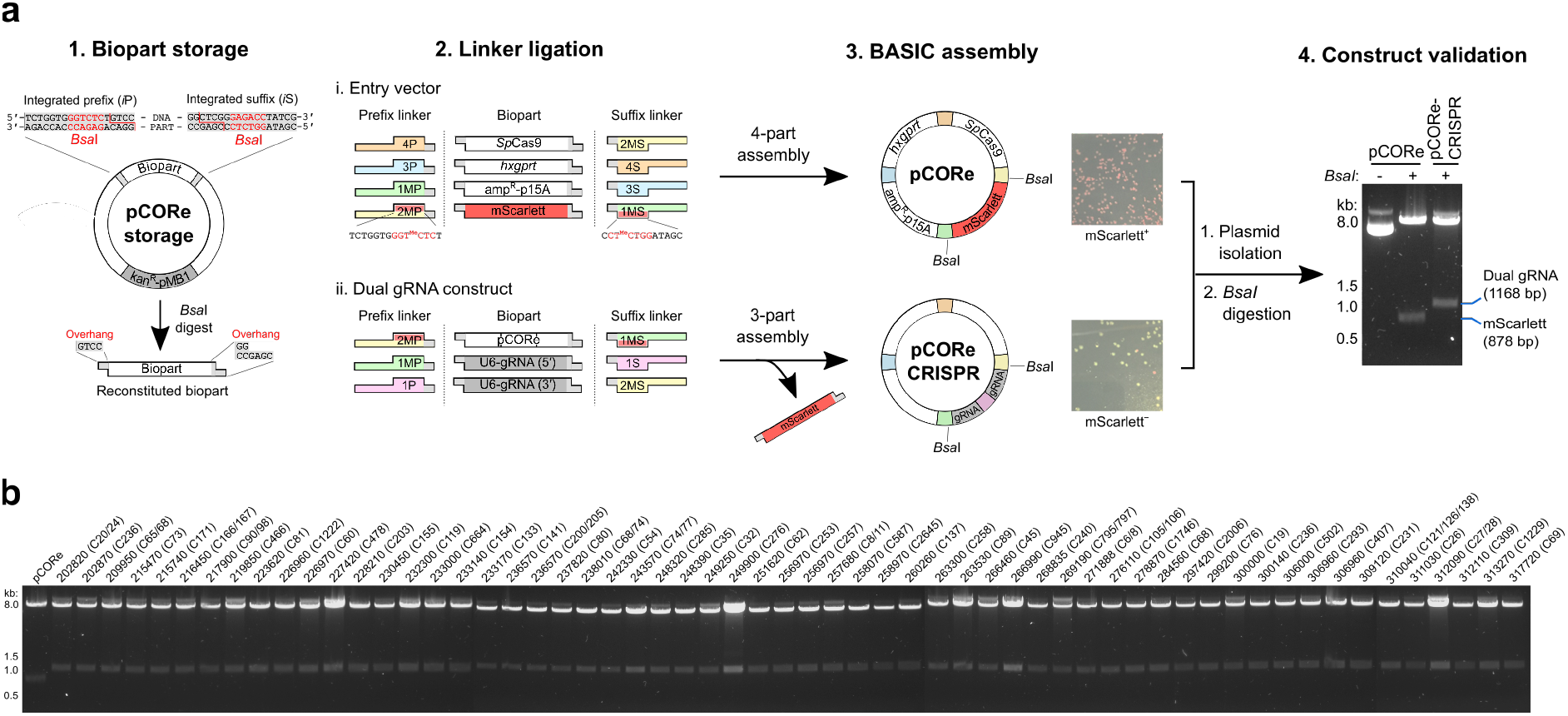
Biopart Assembly Standard for Idempotent Cloning (BASIC) enables modular, high-throughput assembly of CORe CRISPR plasmids. **a.** BASIC strategy used for plasmid construction. (1) The BASIC physical DNA standard. Functional DNA bioparts are flanked by *i*P and *i*S sequences, each containing a *Bsa*I restriction site (red). In CORe, BASIC parts are released from kanamycin-resistant storage plasmids (pCORe storage) by *Bsa*I digestion, enabling the ligation of oligonucleotide linkers for subsequent vector assembly via the BASIC workflow^3^. (2,3) Assembly strategy for the CORe entry vector (‘pCORe’; i) and final dual gRNA constructs (‘pCORe CRISPR’; ii). pCORe is generated through the ordered assembly of four bioparts: *Sp*Cas9, *hxgprt*, amp^R^-p15A and mScarlett. Bacterial transformants of pCORe exhibit a pink phenotype due to expression of the mScarlett fluorophore. The methylated cytosines uniquely present in the linkers flanking mScarlett prevent digestion of the linker during the assembly process and reconstitutes pCORe (*Sp*Cas9-*hxgprt*-amp^R^-p15A) for a second round of assembly. The pCORe biopart is then subject to a 3-part assembly reaction with two gRNA parts, replacing the mScarlett cassette and generating pCORe CRISPR. Transformants of pCORe CRISPR appear non-fluorescent due to the loss of the mScarlett marker, enabling rapid selection of successful assemblies. (4) Following plasmid isolation, successful insertion of the two gRNA parts is verified by differential size analysis of fragments upon *Bsa*I digestion. **b**. *Bsa*I verification of a 59-member pCORe CRISPR library targeting 74 hyperreactive cysteines of *T. gondii*; successful gRNA insertion was achieved for all selected clones.

**Figure S4.**
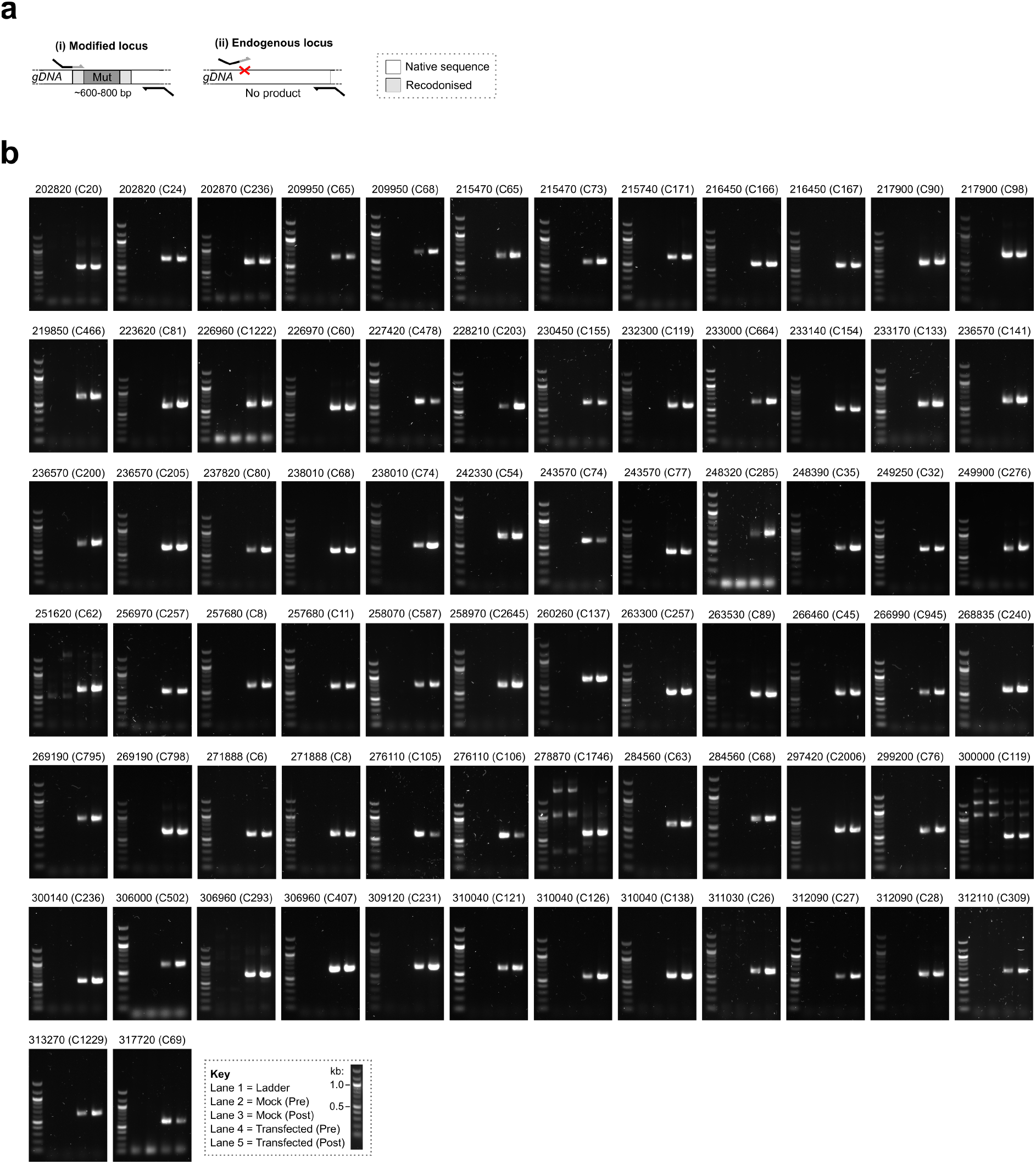
Integration-specific PCR enables selective amplification of CORe mutant DNA across diverse genomic loci. **a.** General PCR strategy for generating amplicons encompassing the modified cysteine loci of mutant parasites. Specific amplification of modified vs. endogenous genomic loci is achieved by priming regions of unique recodonized sequence in the integrated mutational templates. **b.** Integration-specific amplicons for 74 cysteines targeted for mutagenesis via CORe. For all targets, mutants are detected in both ‘Pre’ and ‘Post’ timepoints, and no product is generated in mock-transfected parasite populations.

**Figure S5.**
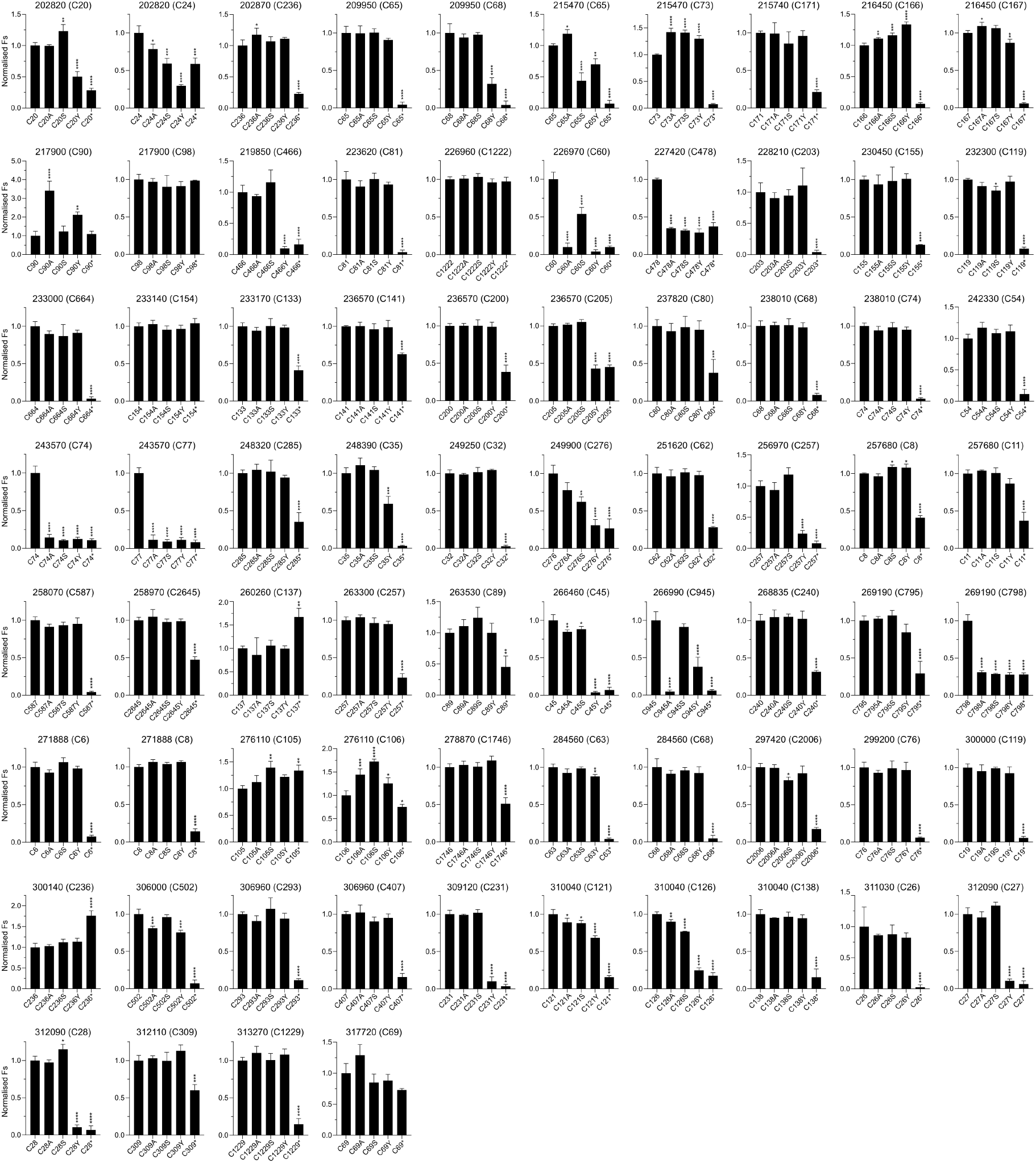
Hyperreactive cysteines in *T. gondii* targeted by CORe exhibit reproducible and diverse mutational profiles. Histograms showing normalized Fs values for five mutations (a recodonized cysteine codon, alanine, serine, tyrosine and stop codon) following mutagenesis of 74 reactive cysteines in *T. gondii* with CORe. The gene identifier (from ToxoDB; https://toxodb.org/) and associated cysteine residues are annotated above each plot and organised numerically in ascending order. Data represents mean ±s.d. for 3 biological replicates (n=3). Statistical significance of the non-synonymous mutations against the recodonized cysteine control was determined by one-way analysis of variance. \*\*\*\**p* < 0.0001; \*\*\**p* < 0.001; \*\**p* < 0.01; \**p* < 0.05.

**Figure S6.**
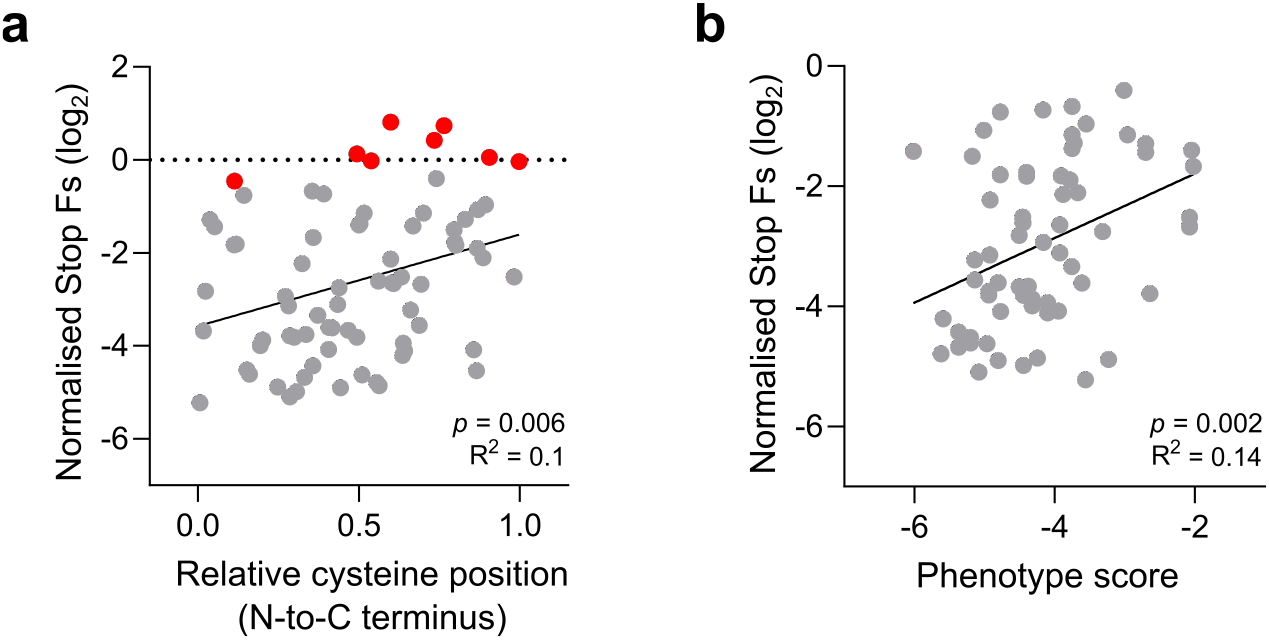
CORe stop codon mutant fitness does not correlate with cysteine position or gene phenotype scores. **a.** Linear regression analysis of normalised stop codon Fs values against the relative position of the mutagenized cysteine in the associated protein sequence. While no overall correlation is observed, targets without a statistically significant stop codon phenotype (coloured red) generally cluster toward the C terminus. **b.** Comparison of significant stop codon mutant phenotypes against published gene phenotype scores^4^; no overall relationship is observed. Annotated R^2^ values indicate the degree of correlation between datasets being compared.

**Figure S7.**
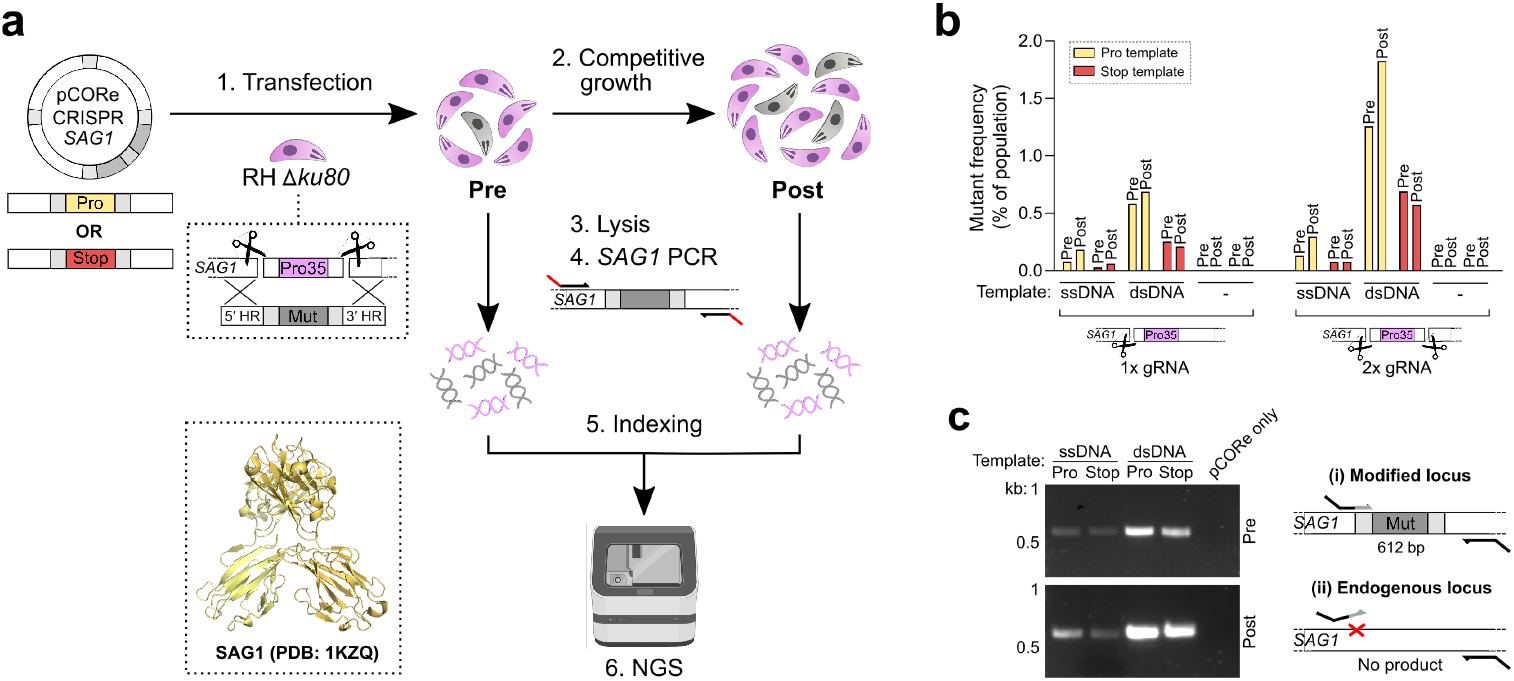
Optimal editing efficiency at *SAG1* is achieved using a dual gRNA strategy with double-stranded template conformation. **a.** Workflow used for assessing the integration efficiency of templates for site-directed mutagenesis of *SAG1* (P35) with a synonymous recodonized proline (wildtype) or stop codon (knockout). **b.** Frequency of *SAG1*-modified parasites before (‘Pre’) and after (‘Post’) a period of competitive lytic growth. Templates were provided in either single (ssDNA) or double-stranded (dsDNA) conformation and transfected with single or dual gRNA-containing pCORe CRISPR plasmids. For each mutation type, a maximum integration frequency (1-2%) is achieved following transfection of dual gRNA plasmids with dsDNA templates. **c.** Template-specific PCR from dual gRNA samples showing selective amplification of proline mutant DNA. Data represents a single experiment (n=1).

